# Single-cell transcriptome reveals a testis-specific expression profile of *TCEA* in human spermatogenesis

**DOI:** 10.1101/2021.03.10.434830

**Authors:** S. Pankaew, P. Pramoj Na Ayutthaya

**Affiliations:** Aix-Marseille Université, CNRS, INSERM, CIML, 13009 Marseille, France; Department of Molecular Biology, Max Planck Institute for Developmental Biology, 72076 Tübingen, Germany

## Abstract

Transcription elongation factor A (*TCEA*) is a eukaryotic transcriptional molecule, required for a formation of initiation and elongation of gene transcription-mediated RNA polymerase II (RNAPII) complex, to promote transcription-coupled nucleotide excision repair (TC-NER) after RNAPII backtracking recovery. *TCEA* shares three isoforms in which *TCEA1* is ubiquitously expressed among all eukaryotic cells. We found a spermatogenesis *TCEA1* and *TCEA2* expression profile has a unique transcriptional programme, compared with embryogenesis. Moreover, the testis-specific *TCEA2* profile correlates with gene transcription, whereas *TCEA1* specifically correlates with genes transcribed for Nuclear excision repair (NER) during human spermatogenesis. We also found that the expression activation of *RNF20*, a *TCEA1* inhibitor, leads to expressional *TCEA1* reduction, but having no direct impact on *TCEA2* expression, implying the potential *RNF20*-dependent transcriptional switching of *TCEA2* in transcriptional regulation during spermatogenesis. Our analysis defined a transcriptional bursting event where transcription-coupled repair (both Base excision repair and Nuclear excision repair) is a major pathway highly expressed in early spermatogenesis, supporting the transcriptional scanning hypothesis of which mutation of transcribed genes is effectively repaired as proposed by Xia B., et al. (2020).

## Introduction

Transcriptional bursting is a fundamental biological process signatured in developments such as murine embryogenesis and spermatogenesis (Li et al., 2013; Ochiai et al., 2020). Upon the transcription, massive transcription factors are expressed to help promote downstream transcribed genes (Rhee et al., 2008). Transcription factor IIS (TFIIS) or transcription elongation factor A (TCEA) is one of the most crucial molecule to coordinate gene transcription via forming an initiation and elongation complex with RNA polymerase II (RNAPII) (Guglielmi et al., 2007; Kim et al., 2007; Prather et al., 2005; Reines et al., 1989) and also to help safeguard transcriptional error-prones from RNAPII complex by possessing a helicase activity that dissociates mismatched transcribed DNA lesion via mRNA cleavage of transcription-coupled nuclear excision repair or TC-NER (Kalogeraki et al., 2005; Kettenberger et al., 2003; Sigurdsson et al., 2010). Among Chordata phylum, *TCEA* shares three isoforms: *TCEA1*, *TCEA2*, and *TCEA3* and only *TCEA1* has been investigated at biochemical level. Previous study demonstrated that the RNAPII complex with TCEA1 can help recover RNAPII backtracking (Lisica et al., 2016) upon promoting a bypass via a UV-induced 8-oxoguanine presence (Kuraoka et al., 2007). Yet it is unclear whether other isoforms share a similar expression profile as *TCEA1* and how those isoforms biochemically regulate downstream pathways. Eukaryotic *TCEA* is also crucial for biological developments across kingdoms ranging from yeast, plant to animal (Cha et al., 2013; Grasser et al., 2009; Guglielmi et al., 2007; Park et al., 2013; Yang et al., 2018). Dysfunction of *TCEA1* significantly leads to cancer proliferation, embryonic lethality, and sperm abnormality in human (Horiuchi et al., 2018; Ito et al., 2006; Shema et al., 2011) due to severely reduced gene transcriptions and genome stability (Lennon et al., 1998; Zatreanu et al., 2019). A recent work from Xia B., et al. (2020) has demonstrated that SNP variants in transcribed genes have significantly lowered mutation rates rather than those in non-transcribed genes, suggesting the gene repairs can be fine-tuned via transcription-coupled repair during gene transcription in spermatogenesis (Xia et al., 2020).

Intriguingly, expression patterns of *TCEA1*, *TCEA2*, and *TCEA3* showed a unique characteristic in *Xenopus* embryogenesis (Labhart and Morgan, 1998), hypothesizing that these three isoforms may have functional preferences for both gene transcription and DNA repair based upon their gradient expressions among each cell stage. *TCEA2* was reported, in 1997 to be testis-specific gene (Weaver and Kane, 1997). But how *TCEA2* is quantitatively expressed at single-cell level is not known. In this work, we sought to investigate if *TCEA1*, *TCEA2*, *and TCEA3* expressions can be correlated with gene transcription and DNA repair expression in human spermatogenesis. Collectively, our study reports the germline-specific *TCEA1 and TCEA2* transcription in which the expressed isoforms can refer to the functional preference of gene transcription and the repair. This analysis would help highlight a new outlook of expression-based *TCEA* isoforms, which infer a new mechanism of *RNF20*-independent *TCEA2* regulation in the context of developmental biology.

## Result and Discussion

### Transcription elongation factor *TCEA1* and *TCEA2* expression profiles correlate with cellular gene transcription during spermatogenesis

A previous study from Xia B., et al. (2020) generated a hypothesis “transcriptional scanning” in which expressed genes can be effectively repaired via a well-known characteristic of transcription-coupled repair (TCR) and that can promote an impact of gene mutations of non-transcribed genes in mammalian testes (Xia et al., 2020). We here investigated what molecular factors contributing to the transcription and the repair can be a characteristic of fixing spermatogenesis-expressed gene mutations at gene transcription level. To test dataset robustness and reproducibility, we performed an independent statistical analysis and identified the specific cell types (Figure 1A) by using gene markers from human testes atlas (Supplementary Table 1). Overall, our statistical analysis provided similar results given by principal components embedding and identified groups of spermatogenesis-related genes which are highly expressed among individual cell stages by comparing with known marker genes (Figure 1B&Supplementary Figure 1A&1B) as similar to the human testes atlas study (Guo et al., 2018). Lists of germline-selective differentially expressed gene clusters are provided in Supplementary Table 1. Human gene transcription is highly expressed in round spermatid (RS) stage with low leakage of mitochondrial gene expression, suggesting the stage was not under stress-induced, but rather cell proliferation- or morphogenesis-induced scenario (Figure 1B&Supplementary Figure 1C&1D). We found that the expression profile of *TCEA1* and *TCEA2* showed a specific pattern (Figure 1C), in which during the early stage of spermatogenesis from Sg-1 to RS-1, *TCEA1* expression decreased as *TCEA2* expression increased. Later, the expression *TCEA1* were recovered during the stage RS-2 to ES-4. As expected, *TCEA* enhances transcription elongation to help increase gene transcription rate in complex with RNAPII (Schweikhard et al., 2014a; Schweikhard et al., 2014b). Interestingly, the gradient pattern of *TCEA1* and *TCEA2* shows a unique characteristic only observed in spermatogenesis not as in developmental and tissue-specific characteristic in embryogenesis from humans (Supplementary Figure 2) and from *Xenopus (Labhart and Morgan, 1998; Sladitschek et al., 2020)*.

**Figure 1.**
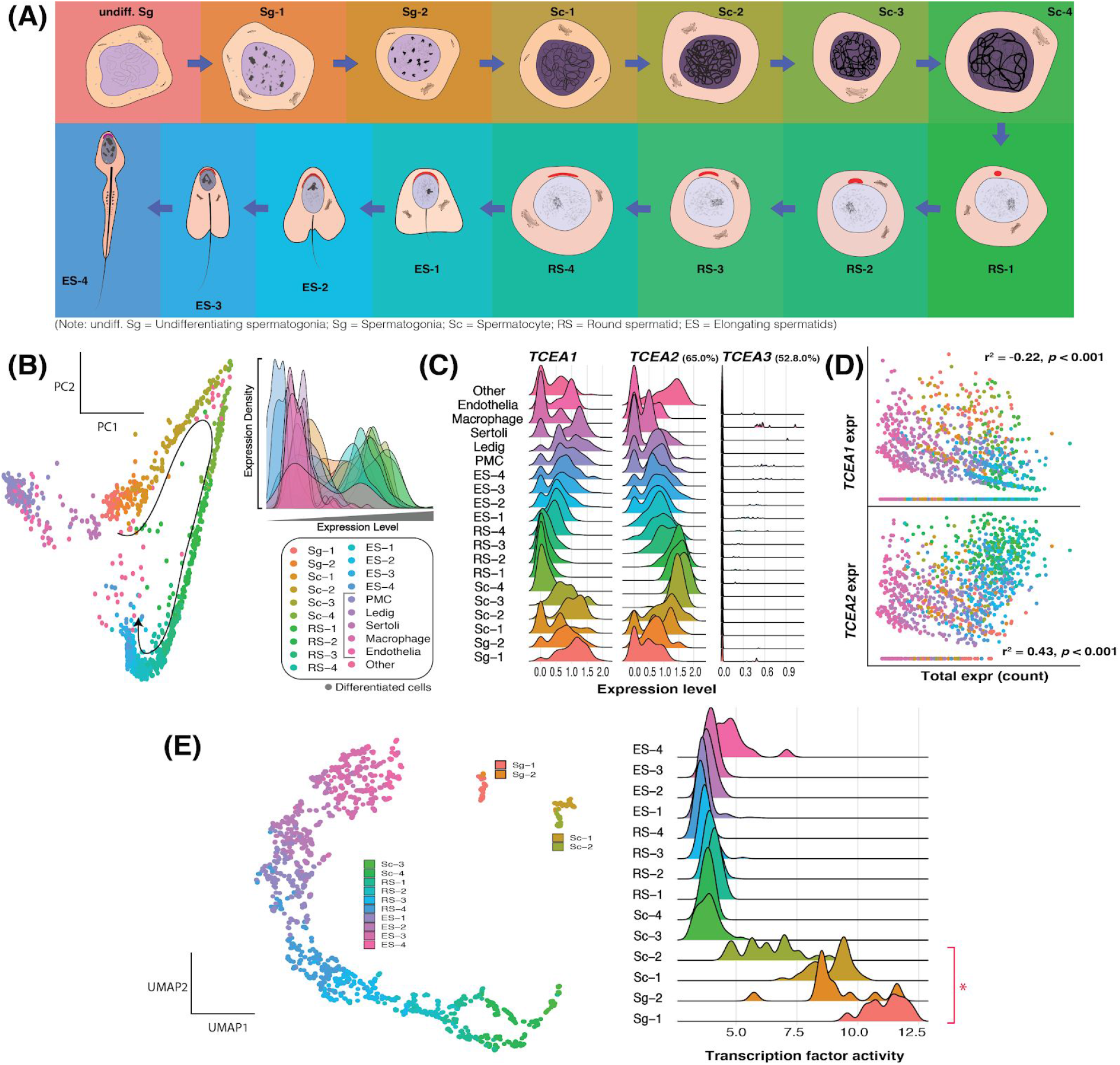
Gradient expression of *TCEA1* and *TCEA2* with cellular gene transcription upon developmental niches of spermatogenesis. (A) Schematic spermatogenesis-related germlines given in the analysis. (B) Principal component analysis (PCA) of scRNA dataset provided by Xia B., et al. (2020) shows cell type-specific trajectories upon gene transcription in spermatogenesis with total gene transcription profiles of each cell type in humans. (C) Transcriptional profiling of human *TCEA1*, *TCEA2* and *TCEA3* given protein sequence identity 65.0% of TCEA2 and 52.8% of TCEA3, compared with TCEA1 in individual cell types. (D) *TCEA1* and *TCEA2* expression count correlates statistically with gene transcription in spermatogonial-specific germlines. (E) Profiling of transcription factor activity in human spermatogenesis (high activity from Sg-1 to Sc-2 as indicated with red asterisk) provides unique characteristics of transcription factors involved in the spermatogenesis given by uniform manifold approximation and projection (UMAP, left) and quantitative activity profile (right). **[all statistical analysis provided in the figure was tested by Pearson’s correlation at a cut-off *p*-value = 0.001]**

*TCEA* is also contributed to help balance ribosomal gene transcription under cell stress (Gomez-Herreros and de Miguel-Jimenez, 2012), but that is not likely our observation because mitochondrial gene leakage as inferred to cell stress is low during spermatogenesis (Supplementary Figure 1D). Next, we found that gradient total gene transcription of each stage ranging from spermatogonia (Sg) to round spermatid (from low to high) correlates negatively with *TCEA1* and positively with *TCEA2*, but not as similar case in human embryogenesis (Figure 1D&Supplementary Figure 2). Therefore, the expression profile of *TCEA1 and TCEA2* can be unique in human spermatogenesis. Based on assumptions: (i) total encoding gene being transcribed is mediated by RNAPII in complex with TCEA, and (ii) each isoform of *TCEA* can be translated into functional TCEA. We reasoned that the molecular testis-specific *TCEA2* preference after translation can promote transcription regulations due to “transcriptional *TCEA* switching” over other isoforms for spermatogenesis-related gene transcription homeostasis. To investigate transcriptional activity profiling during the spermatogenesis, we embedded the uniform manifold approximation and projection of the transcription factor expressions of which the relevant genes shared their TF motifs are given high expression scores to infer the TF or regulon activity. We observed high regulon activities and unique trajectories during the early spermatogenesis (from Sg-1 to Sc-2), compared with the rest of cell stages that share the continuous trajectories (Figure 1E). That suggests that the early spermatogenesis has a unique transcriptional mechanism and regulation to maintain important gene expressions prior to initialising round and elongating spermatid differentiation.

### Nuclear excision repair is a major DNA repair expressed during human spermatogenesis

We calculated regulon activities using Single-Cell rEgulatory Network Inference and Clustering (SCENIC) (Van de Sande et al., 2020), a computational method for inferring gene regulatory network and its activities. The inferences calculated co-expression between a transcription factor and its targets, clustered them into a ‘regulon’, and computed the regulon activities from the target genes in single-cell. We found high regulon activities in the early spermatogenesis (from Sg-1 to Sc-3) with high number of regulon interactions, compared to other cell stages (Figure 1E&Figure 2A). To investigate downstream gene functions regulated by the relevant regulons, we performed gene ontology (GO) analyses with an adjusted *p*-value cutoff at 0.01. List of the regulons and downstream genes identified in our analyses is provided in Supplementary Table 2&Supplementary file 1. Both positive and negative regulations of RNAPII-mediated transcription were shown prominent during the Spermatogonia and Spermatocyte (only Sc-1 and Sc-2), while Sc-3 has an unclassified regulation of the transcription (Figure 2B). However, positive regulation of the transcription regulation was not found in other cell stages (Supplementary Figure 3). This suggests that high positive activity of the RNAPII-mediated gene transcription, so called “transcriptional bursting” occurs in the specific early stages of the spermatogenesis from Sg-1 to Sc-3. Given the transcriptional scanning hypothesis, transcriptional bursting may promote gene repairs to fine-tune gene adaptations due to high transcription rate and transcription-coupled repair expression (Svejstrup, 2002; Xia et al., 2020). We therefore performed correlation analyses of the regulon activity and each of the DNA repair pathways. We found that both Base excision repair (BER) and Nuclear excision repair (NER) show high correlations with the regulon activity, suggesting that preferential transcription-coupled repair mediates transcribed gene repairs during the transcriptional bursting (Figure 2C&Supplementary Figure 6A). In post-transcriptional bursting, the NER pathway showed the highest expression among other DNA repair pathways (Figure 2D&Supplementary Figure 6B).

**Figure 2.**
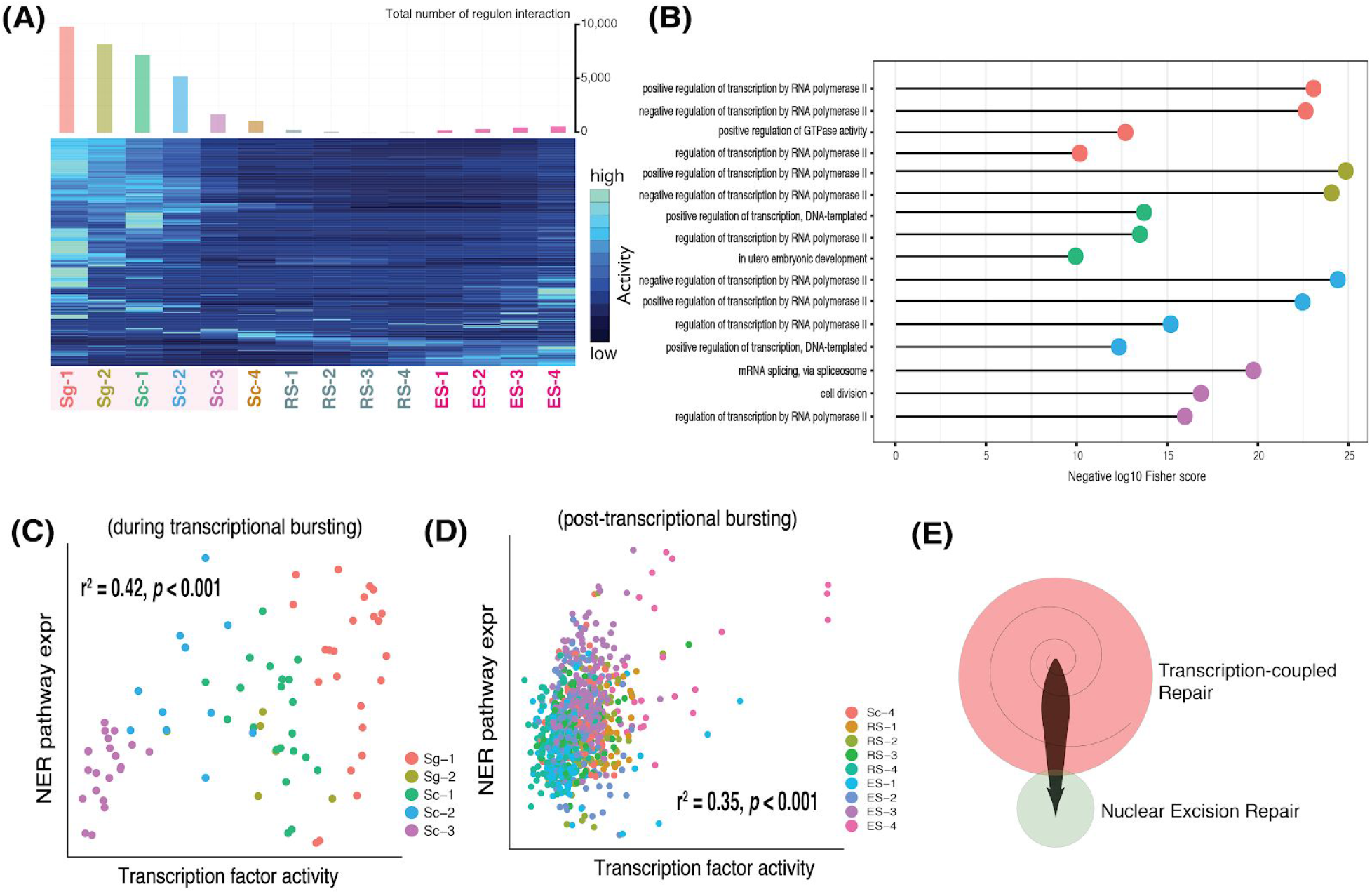
Defining transcriptional bursting during human spermatogenesis, which shares a high correlation with nuclear excision repair (NER) expression. (A) Heatmap representation of regulon activity profiling of each cell stage during spermatogenesis with a number of regulon-targeted gene networks represented in the histogram. (B) Selected gene ontology (GO) analysis of genes highly regulated by regulons involved in RNAPII-mediated gene transcription in Sg-1, Sg-2, Sc-1, Sc-2, and Sc-3 represented by colours given from the heatmap. (C) Correlation analysis of NER pathway expression with transcriptional bursting activity by the pySCENIC. (D) Correlation analysis of NER pathway expression with transcription activity after the bursting by the pySCENIC. (E) Schematic model represents NER as a major DNA repair facilitating transcribed gene repair during the transcriptional scanning (both in the transcriptional bursting; red, and the post-transcriptional bursting; light green) as proposed by Xia B., et al. (2020). **[correlation analyses were tested by Pearson’s correlation with a cut-off *p*-value = 0.001; gene ontology (GO) analyses were tested by Fisher’s exact test with a cut-off *p*-value = 0.01]**

The analysis here indicated an important biological regulation upon the transcriptional scanning scenario. Transcriptional-coupled repair is co-expressed with the coupling transcription activity during the early cell stage (Sg-1 to Sc-3) of the human spermatogenesis, whereas NER is primarily co-expressed during the post-transcriptional bursting as shown in the hypothetical model (Figure 2E). This transcriptional bursting event is likely cell-stage specific regulation during the human germline development and highly involved with the transcriptional scanning event because the transcription-coupled repair is responsible for reducing gene mutations during which high activities of RNAPII-mediated gene transcription. Overall, the transcriptional bursting in the early spermatogenesis is potentially responsible for the transcriptional scanning hypothesis in which the transcription-coupled repair plays a major role in reducing mutations of transcribed genes and promoting gene adaptations.

### *TCEA1* expression infers preferential transcription involved in transcription-coupled repair in human spermatogenesis

We next investigated *TCEA1* and *TCEA2* expression, which may transcriptionally correlate with the spermatogenesis-related NER expression. We realized that having a single marker to determine each of the DNA repair pathways would introduce less plausible pathway expressions due to high variants of single-cell expression counts. Therefore, we curated gene lists grouped into individual gene clusters, representing unique DNA repair characteristics via our knowledge-based approach. List of genes involved in the DNA repair pathways is provided in Supplementary Table 3. Albeit nuclear excision repair (NER) is biochemically classified into two pathways: TC-NER and general genome NER (GG-NER), it is difficult to interpret whether TC-NER is more highly expressed than GG-NER given in our dataset without protein activity. We then created the NER-related gene cluster to infer TC-NER/GG-NER expression, which is highly expressed in Sc stage, with the presence of specific TC-NER markers such as *POLRA2*, *ERCC6* or *ERCC8* and *XPC*, respectively (Supplementary Figure 4A,4B,4C,4D and Supplementary Figure 5). We found that only *TCEA1* has the highest correlation with NER pathway, but not in the case of *TCEA2*. We also plotted *TCEA1* and *TCEA2* against other DNA repair expression profiles (Figure 3A). Although all DNA repair pathways were highly expressed during the early spermatogenesis, neither *TCEA1* nor *TCEA2* showed strong correlations with Base excision repair (BER) and Mismatch repair (MMR) (Supplementary Figure 4E&4F&Supplementary Figure 5), implying only *TCEA1*-mediated TC-NER during gene transcription.

**Figure 3.**
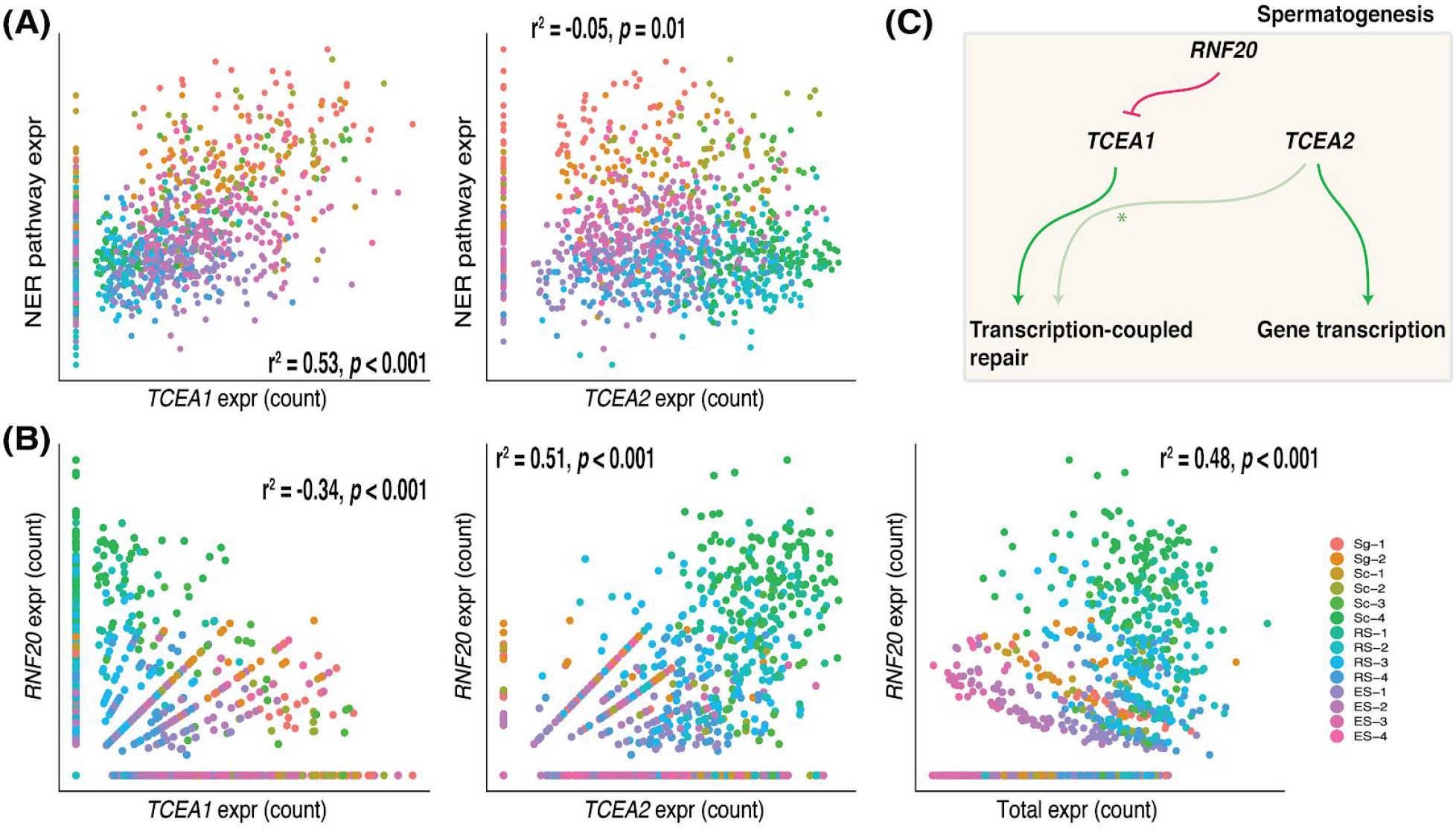
Comparative expression correlations between *TCEA1* or *TCEA2* and DNA repair pathways and analysis of DNA repair pathway expression in human spermatogenesis. (A) Correlation analyses of NER pathway expression with *TCEA1* and *TCEA2*. (B) Correlation analyses of *RNF20* regulations upon *TCEA1* (left), *TCEA2* (middle) expression and gene transcription (right). (C) Proposed gene interaction of *RNF20*-mediated *TCEA1* and preferential *TCEA* switching based on transcription profiling [preference, green; inhibition, red; no correlation, asterisked light green]. **[all statistical analysis provided in the figure was tested by Pearson’s correlation at a cut-off *p*-value = 0.001]**

Messenger RNA from gene transcription can be implemented to facilitate gene repairs via NHEJ mechanism during gene transcription (Chakraborty et al., 2016). That would also speculate that mRNA products can be utilized to repair transcribed genes via NHEJ as we observed a positive correlation between *TCEA2* and NHEJ pathway with high cellular gene expressions (Supplementary Figure 1C&4F). We also found that NER pathway is highly expressed given high expressions among *POLR2A*, *ERCC6*, and *ERCC8*, but low *XPC* expression, implying the correlation of *TCEA1* expression infers the TC-NER pathway activity (Figure 3A&Supplementary Figure 4). Moreover, a previous report demonstrated that *ERCC6*, which encodes Cockyne Syndrome B (CSB), crucially insures a preferential DNA repair during the G2/M-phase of cell division, leveraging transcription-coupled related homologous recombination or proposed TC-HR in transcriptionally active chromatin (Aymard et al., 2014; Batenburg et al., 2015). In order for “pushing” a stalled CSB-RNAPII complex, *TCEA* is required to recover the RNAPII-mediated transcription elongation. Speculatively therefore, TC-HR would need a required amount of *TCEA* to facilitate the repair as we observed the positive correlation between *TCEA1* and HR pathway during spermatogenesis (Supplementary Figure 4E). *TCEA1* expressed during Homologous recombination (HR) while *TCEA2* expressed during Non-homologous end-joining (NHEJ) and Inter-crosslinking repair (ICL), which also highly expressed and involved in S-phase (Supplementary Figure 7) in which DNA replication and ICL repair is a crucial prerequisite to fix mutagenic lesions at a replication fork for the cell cycle progression in the early and mid stage (Sg to RS) of spermatogenesis (Supplementary Figure 4&5). Given a high scoring expression of spermatogenesis G2/M-phase during cell cycle (Supplementary Figure 7), we found that cell division-relevant DNA repairs such as HR and NHEJ were highly cooperated during the early spermatogenesis, but more often expressed for only NHEJ during RS stage (Supplementary Figure 4&6) because NHEJ is more efficient and faster to facilitate DSB repair, but less accurate than HR for genomic stability (Bee et al., 2013; Derbyshire et al., 1994; Mao et al., 2008; Shrivastav et al., 2008). We detected an increased expression of *RNF20*, a known inhibitor of *TCEA1* from the stages Sg-3 to RS-3 (Supplementary Figure 1E). We also plotted *TCEA1* and *TCEA2* against *RNF20*, a crucial inhibitor suppressing TFIIS activity in human cells for transcriptional *TCEA* balance. We found that *TCEA1* has an antagonistic correlation, whereas *TCEA2* has a synergistic correlation with *RNF20* profile during spermatogenesis (Figure 3B). Interestingly, *RNF20* positively correlates with spermatogenesis-related gene transcriptions. This implies that *RNF20* selectively suppresses *TCEA1* activity, but has no direct inhibition towards *TCEA2* gene transcriptions during spermiogenesis (Sc-3 to ES-2). Thus, we proposed a hypothetical model in Figure 3C. Nonetheless, potential molecular suppression related to *TCEA2* has to be further investigated.

### Conclusion

Transcription elongation factor A, consisting encoded three isoforms of which the first isoform *TCEA1* plays crucial roles in transcription initiation and elongation, helps facilitate RNAPII-mediated gene transcription and repair. Despite well-studied *TCEA1* functions, its isoform such as *TCEA2* had been poorly investigated since 1997. Here, we investigated spermatogenesis-related expressional characteristics of individual *TCEA* isoform to help predict what potential molecular preferences would be whether involved in gene transcriptions and repairs by using a single-cell RNAseq data from Xia et al. (2020). We further employed a gene regulatory network inference method (pySCENIC) to infer regulon activities during which the high presence of regulon activity so called “transcriptional bursting” occurred in the early spermatogenesis (Sg-1 to Sc-3). We discovered that transcription-coupled repair expression (both NER and BER) is preferentially co-occurred during the transcriptional bursting, while NER is solely expressed in the post-bursting among other repair pathways. *TCEA1* is shown coexpressing with the NER pathway expression, known for having a helicase activity to facilitate RNAPII backtracking recovery. On the other hand, *TCEA2*, despite sharing a high protein sequence identity, has not been described to share NER regulation.

Interestingly, we showed that *RNF20*, the *TCEA1* inhibitor increases during the post-transcriptional bursting, resulting in the decrease of *TCEA1* expression, while *TCEA2* expression increases along with total gene transcription. With the specific expression profile of testis-specific *TCEA* and its correlation analyses, we propose a hypothetical model of the expressional isoform switching of which *TCEA1* can be down-regulated by *RNF20* after the transcriptional bursting stages, while allow *TCEA2* to facilitate gene transcriptions during human spermatogenesis. Finally, our analysis also supports the transcriptional scanning hypothesis that transcription-coupled repair is highly expressed to help reduce gene mutations during the transcriptional bursting.

## Methods

### Single-cell dataset and analyses using Seurat

Single-cell RNA-Seq and metadata were obtained from Xia B., et al. (2020). The data has already been integrated using Seurat (v3) integration process (Stuart et al., 2019). We performed data quality control by removing cells with less than 200 features expressed and features with less than 3 cells expressing total mRNA transcripts. We also filtered out cells with a percentage of mitochondrial genes > 10%, which represents cells undergoing apoptosis (Osorio and Cai, 2020).

### Dimension reduction plot from principal components

The integrated UMI counts were logarithmically normalized by using NormalizeData function in Seurat and scaled by multiplying by the scale factor of 10,000. From the normalized dataset, 2,000 highly variable expressing genes were selected by using a ‘variance stabilizing transformation’ (vst) from Seurat FindVariableFeatures. These features were then used to perform a principal components analysis (PCA) with all parameters in default. The first two principal components were then used to plot the embedded cells in 2-dimensional space.

### Pathway expression scoring from knowledge-based gene curation

Gene sets involved in DNA repair pathways were manually curated from numbers of literature. We categorized these lists into 6 different pathways containing: Nuclear excision repair (NER), Base excision repair (BER), Mismatch repair (MMR), Homologous recombination (HR), Non-homologous end-joining (NHEJ), and Inter-crosslinking repair (ICL). The list of genes are provided in the Supplementary Table 2. The list of genes in each pathway were then grouped to calculate module expression score, through AddModuleScore in Seurat function (v3) (Stuart et al., 2019).

### Calculating features correlation

We selected Pearson’s correlation to calculate a simple linear correlation between two features. Selected features which are genes, UMI count was used for the calculation. These two selected features were plotted using the FeatureScatterPlot function from Seurat. The statistical analyses and *p*-values were calculated using cor.test function from stats, R default package.

### Inferring transcription factor activity via pySCENIC

We performed regulon activity inference using command line implementation of pySCENIC docker (0.11.0) (Van de Sande et al., 2020). The smoothed expression matrix was used as an input. We computed co-expresssion networks between identified transcription factors and the rest of the genes using default options of pyscenic -grn command. The list of transcription factors were provided in Supplementary Table 3. The list of co-expressed genes were then used to identify enriched DNA binding motifs of transcription factors, using the pyscenic -ctx command. The enriched DNA binding was then used to confirm the presences of a transcription factor and the target genes, and filtered out the target genes, which does not have the motif surrounding its transcription factor start sites (TSS). We used the motif database from motifs-v9-nr.mgi-m0.001-o0.0 and cis-target database mm9-tss-centered-10 kb-7species.mc9nr., both provided by cistarget [https://resources.aertslab.org/cistarget/databases/homo_sapiens]. The resulting co-expression of TF-target is then grouped into regulons. The activity of the regulons were computed using AUCells, which calculates the enrichment of a selected gene set according to the ranking to gene expression in a single cell. First, genes in each cell were ranked according to their expression profile. Then, this rank was used to plot a recovery curve of all target genes in the regulon, and then to compute the ‘Area Under the Curve’ (AUC) as inferred to the regulon activity. We employed the default threshold to compute the area under the curve to top 5 percent of the number of genes in the ranking. The resulting matrix was then integrated into the Seurat object. Finally, using all regulon activities calculated from pySCENIC, we performed a dimension reduction method, using Uniform Manifold Approximation and Projection (UMAP) without scaling and centering. This nonlinear dimension reduction method embedded cells in a 2-dimensional space.

### Regulon analysis

We identified the regulon markers at each stage, using the FindAllMarker function in Seurat. The regulon activity overexpressed in each stage were tested Wilcoxon rank sum test versus the regulon activity from all other stages. To limit the number of significant markers, the threshold of log fold-change was set at 0.1, and adjusted *p*-value (with Bonferroni correction) at 0.01. The list of regulon markers in each stage are in Supplementary Table 3. We then compiled target genes list from stages regulon marker to perform Gene Ontology Enrichment analysis through topGO package. The list of target genes markers were tested with a list of genes in GO terms through Fisher’s exact test. The resulting GO terms were ranked according to Fisher’s test *p*-value.

## Supporting information

regulons.gmt

## Code availability

All codes, docker, expression matrix used in the analysis of this report are available at http://www.github.com/PankaewSaran/PSPP_project

## Author contributions

SP and PP planned the study. SP and PP led data curation and performed the data analysis. SP and PP wrote the manuscript.

## Acknowledgements

We would like to thank Bo Xia and Dr. Itai Yanai for providing the scRNA expression dataset with metadata for our study.

