## Supplementary Information for "Single-cell transcriptome reveals a testis-specific expression profile of *TCEA* in human spermatogenesis"

| | |
|---|---|
| Supplementary Figure 1 | 2 |
| Supplementary Figure 2 | 3 |
| Supplementary Figure 3 | 4 |
| Supplementary Figure 4 | 5 |
| Supplementary Figure 5 | 6 |
| Supplementary Figure 6 | 7 |
| Supplementary Figure 7 | 8 |
| Supplementary Table 1 | 9 |
| Supplementary Table 2 | 23 |
| Supplementary Table 3 | 25 |
| Supplementary References | 26 |

### Supplementary Figure 1

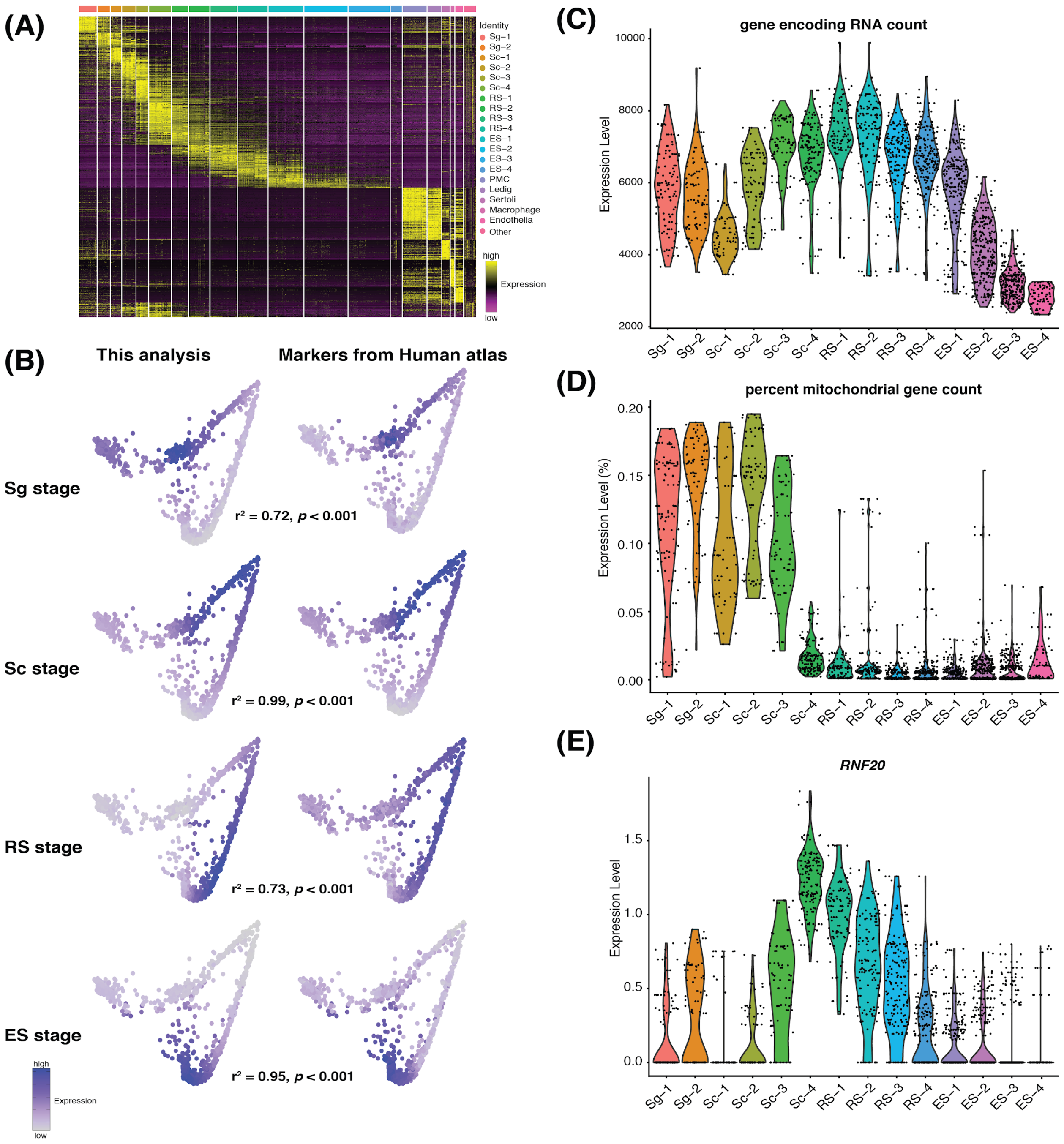

**Supplementary Figure 1. Testing cell-specific dataset by differential expression analysis and known marker genes for gene transcriptions.** (A) Differential gene expression analysis of human spermatogenesis dataset provided by Xia B., et al. (2020) shows unique expression sets among individual cell stages defined by binary logarithmic Fold Change (log2FC) of gene expression at a threshold of 0.25. (B) PCA-based trajectory analysis is consistent with gene sets for cell-stage identifications as reported in human single-cell atlas (Guo et al., 2018). List of gene marker clusters in individual stages is provided in Supplementary Table 1. (C,D,E) Normalized absolute spermatogenesis transcriptome shows total cellular gene transcription, mitochondrial gene transcription, and *RNF20* expression known for *TCEA* inhibition. **[all statistical analysis provided in the figure was tested by Pearson's correlation at a cut-off *p*-value < 0.001]**

### Supplementary Figure 2

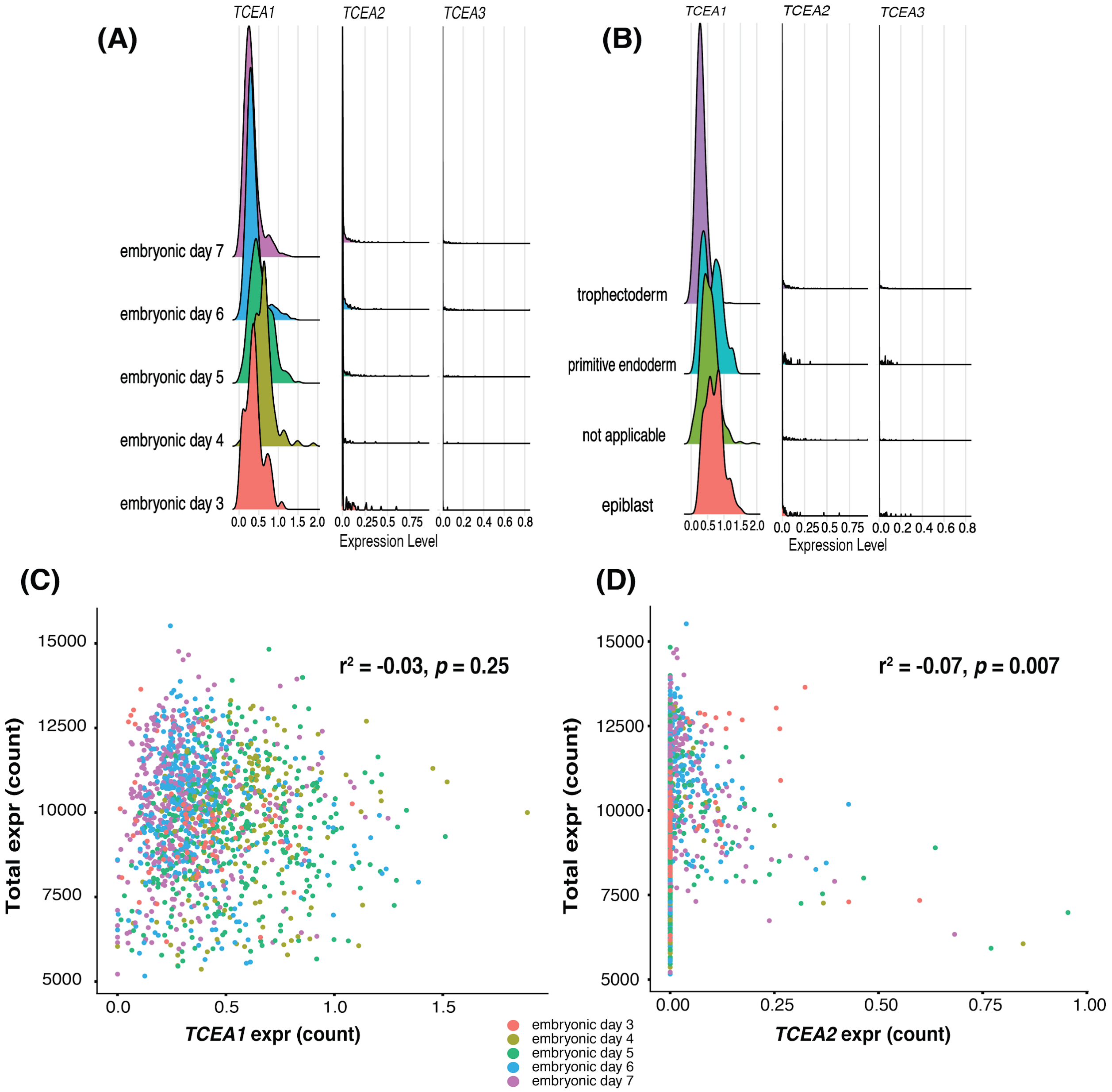

**Supplementary Figure 2. *TCEA* expression profile in human embryogenesis.** (A,B) *TCEA* expression profile of human embryogenesis classified by days [day 3-7] (left) and tissue types [epiblast, not applicable, primitive endoderm, and trophectoderm] (right) provided by (Sladitschek et al., 2020). (C,D) Correlation analyses between embryogene-related gene transcription and *TCEA1* (left) and *TCEA2* (right). **[all statistical analysis provided in the figure was tested by Pearson's correlation at a cut-off *p*-value < 0.001]**

### Supplementary Figure 3

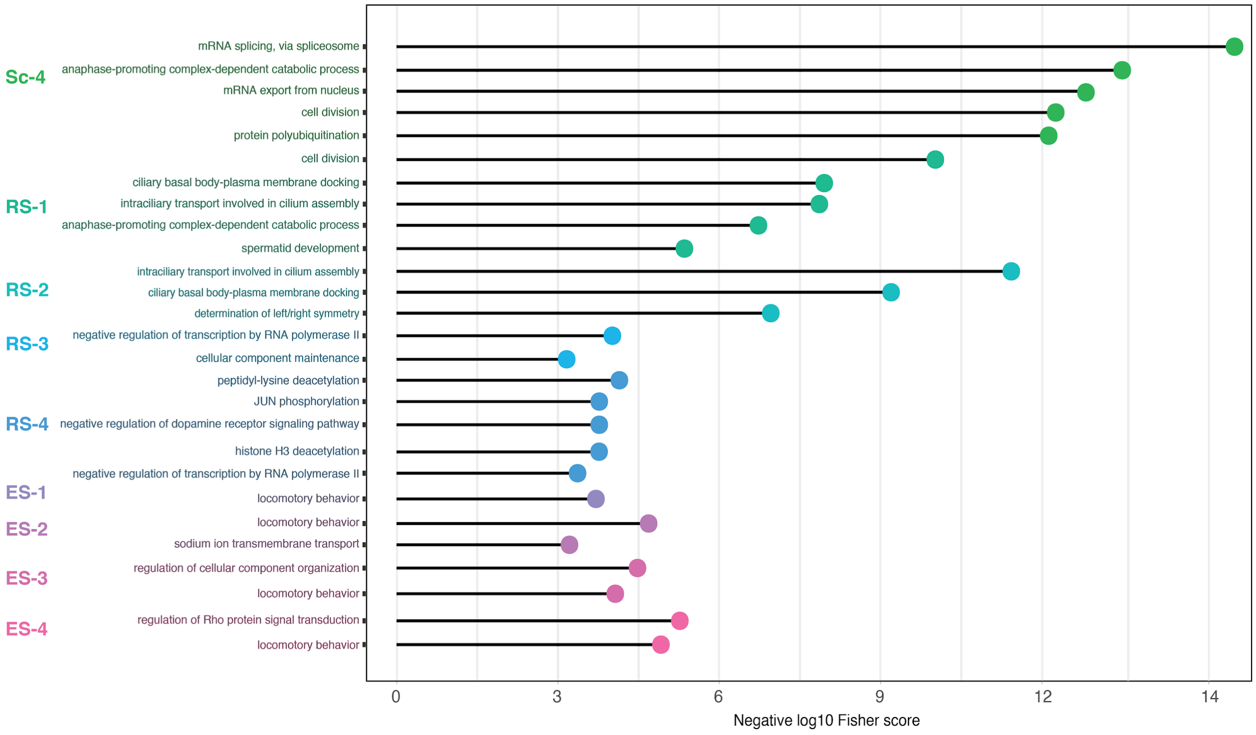

**Supplementary Figure 3. Gene ontology (GO) analysis of genes highly regulated by regulons involved in RNAPII-mediated gene transcription in Sc-4, RS-1 to RS-4 and ES-1 to ES-4.** Each abbreviation provided in the figure is Sc = Spermatocyte; RS = Round spermatid; ES = Elongating spermatid from individual developmental stage. **[gene ontology (GO) analyses were tested by Fisher's exact test with a cut-off *p*-value = 0.01]**

### Supplementary Figure 4

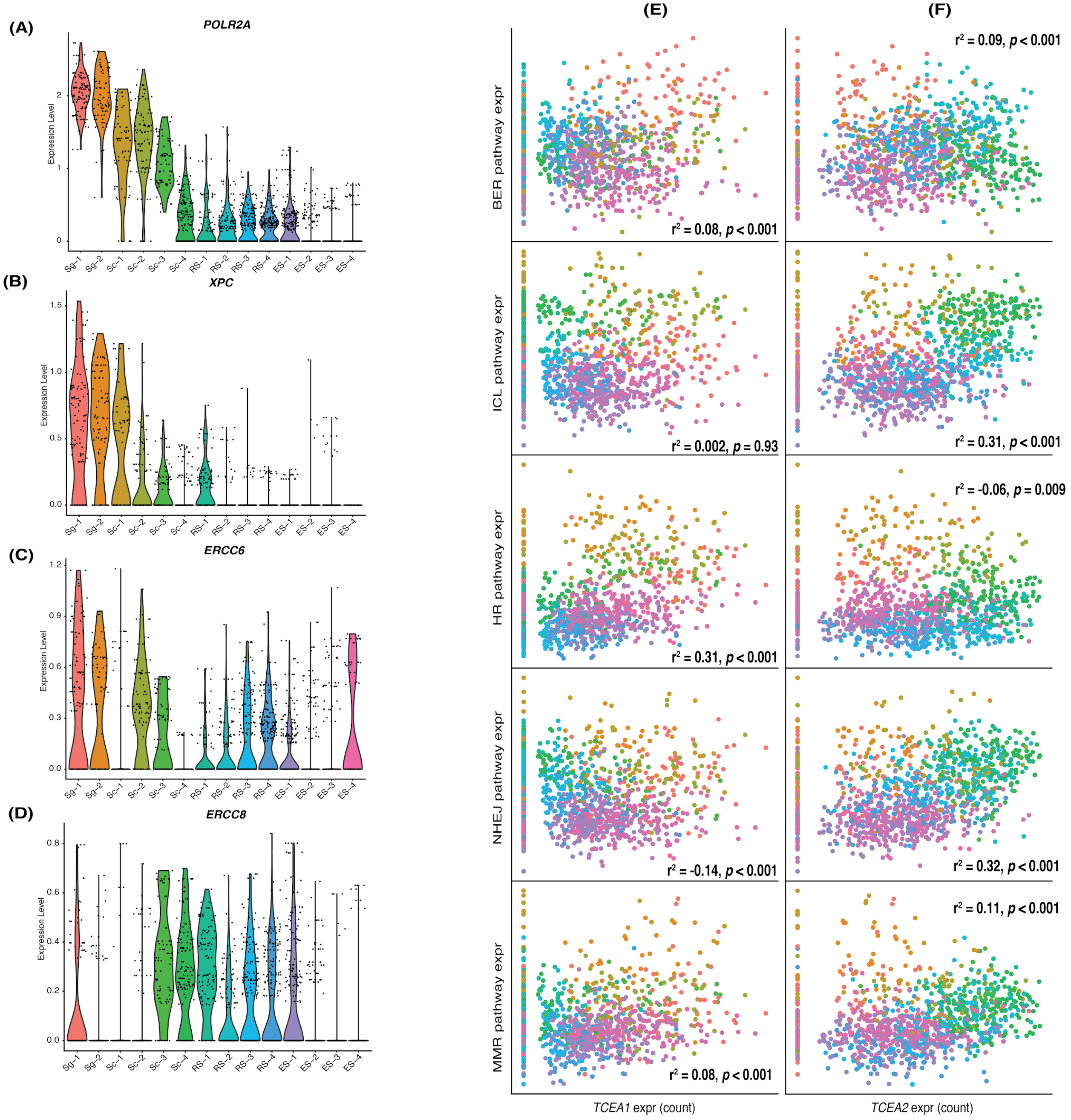

**Supplementary Figure 4. Expression profiles of genes involved in transcription-coupled nuclear excision repair and correlation analyses of *TCEA* cotranscriptionally expressed during DNA repair pathways.** (A,B,C,D) absolute expression profiles of genes involved in TC-NER in individual spermatogenesis-related cell-stages (*POLR2A*, *XPC*, *ERCC6*, and *ERCC8*, respectively). (E,F) Correlation analysis of DNA repair pathway expressions with *TCEA1* and *TCEA2*. DNA repair pathway abbreviation: BER = Base excision repair, MMR = Mismatch repair, HR = Homologous recombination, NHEJ = Non-homologous end joining, and ICL = Inter-crosslinking repair. **[all statistical analysis provided in the figure was tested by Pearson's correlation at a cut-off *p*-value < 0.001]**

### Supplementary Figure 5

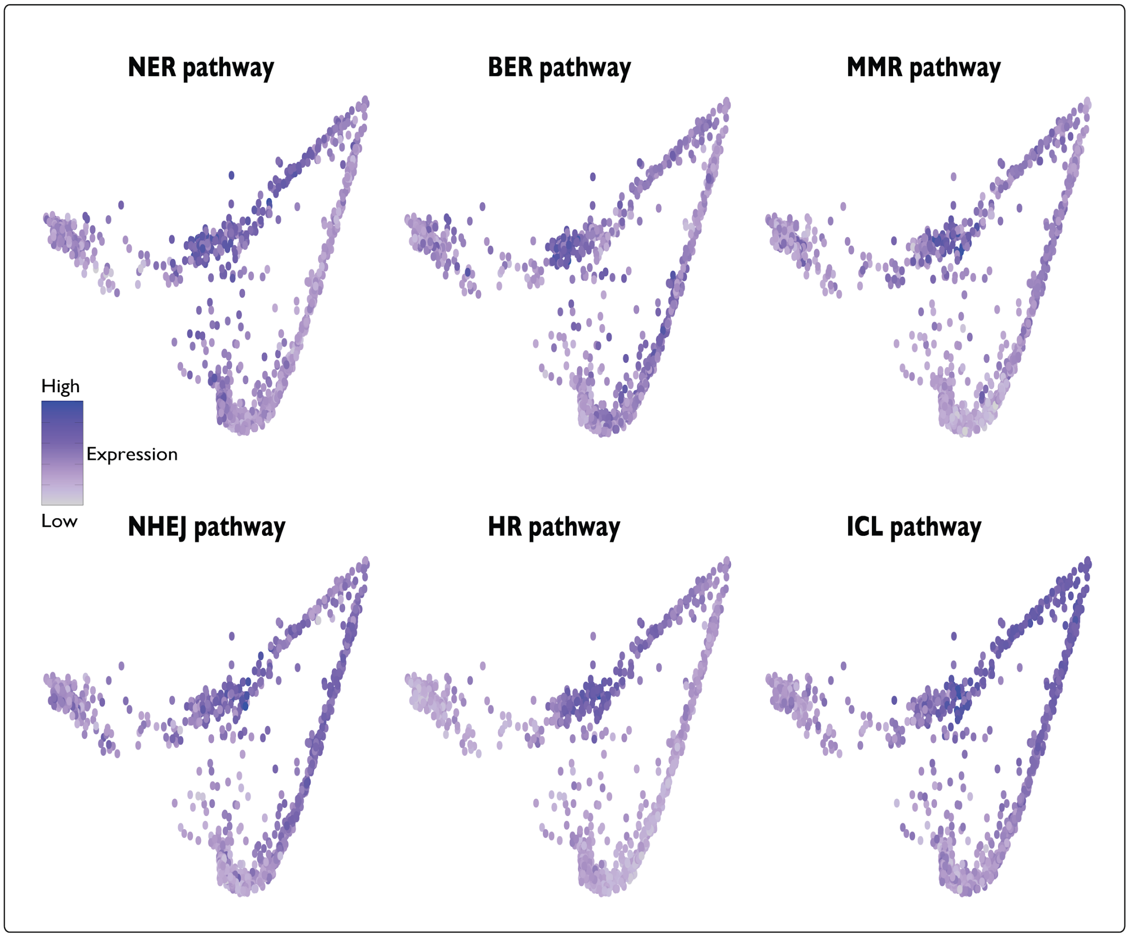

**Supplementary Figure 5. DNA repair pathway expressions in Principal component analysis trajectories in cell-type specific spermatogenesis.** Each of the DNA repairs show high expression in the early stage of Spermatogonia (Sg) during the transcriptional bursting. NHEJ also shows high expression in Round spermatid (RS) stage, and ICL shows high expression in Spermatocyte (Sc) stage as they are responsible for double strand break (DSB) repair in G2/M-phase and S-phase of cell cycle, respectively as we observed in Supplementary Figure 5. Color represents expression level from high (purple) to low (grey). DNA repair pathway abbreviation: BER = Base excision repair, MMR = Mismatch repair, HR = Homologous recombination, NHEJ = Non-homologous end joining, and ICL = Inter-crosslinking repair. List of genes involved in the DNA repair pathways is provided in the Supplementary Table 2.

### Supplementary Figure 6

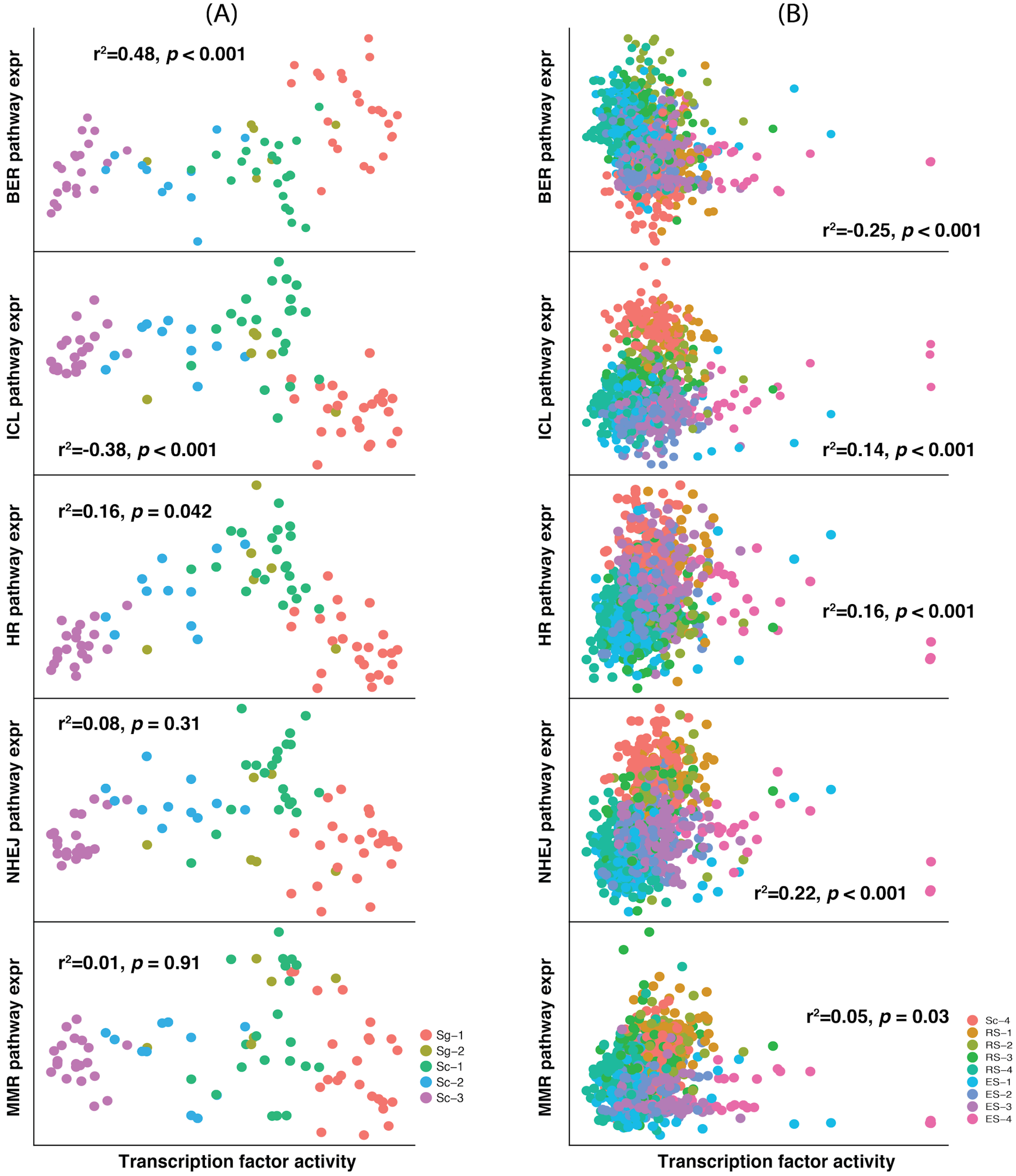

**Supplementary Figure 6. Correlation analyses of DNA repair pathway expression with transcription factor activity defined during the transcriptional bursting (A) and the post bursting (B).** DNA repair pathway abbreviation: BER = Base excision repair, MMR = Mismatch repair, HR = Homologous recombination, NHEJ = Non-homologous end joining, and ICL = Inter-crosslinking repair. **[all statistical analysis provided in the figure was tested by Pearson's correlation at a cut-off *p*-value < 0.001]**

### Supplementary Figure 7

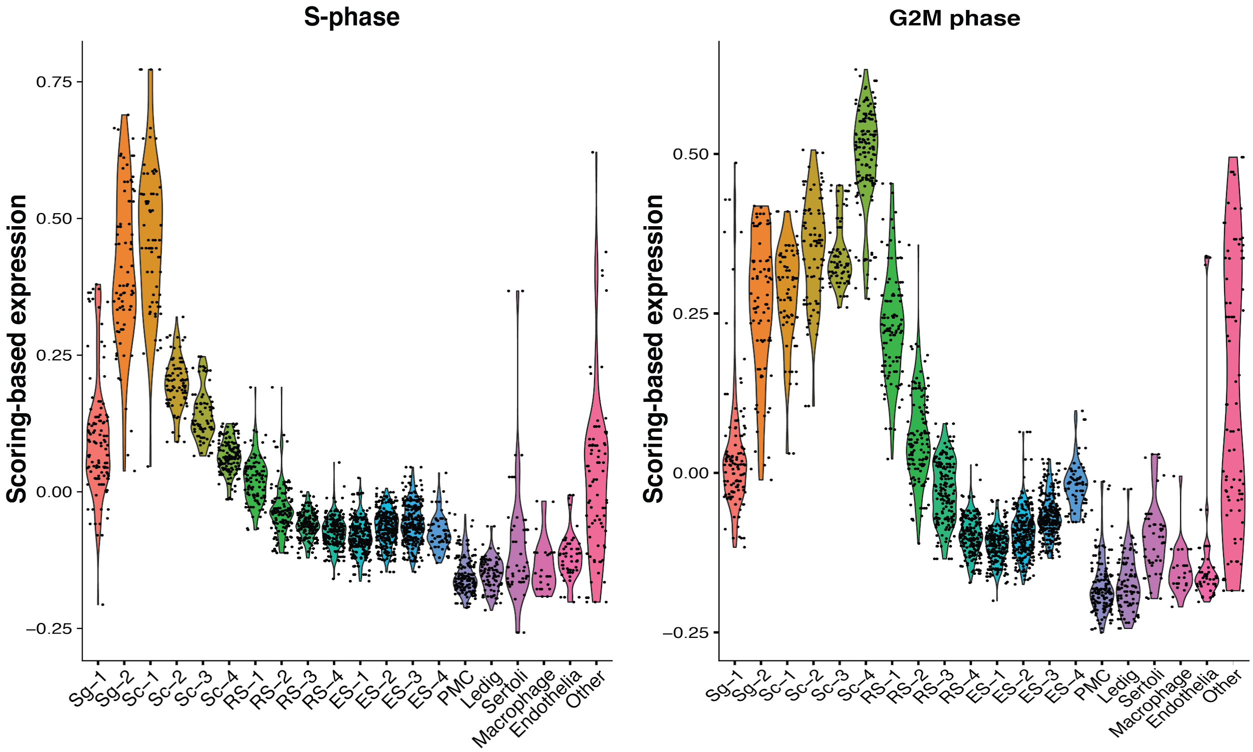

**Supplementary Figure 7. Gene expressions for determining Cell cycle in S- and G2/M-phase during spermatogenesis.** Relative expressions of S-phase (left) and G2/M-phase (right) in the cell cycle of individual cell-type spermatogenesis in human.

### Supplementary Table 1

**Table 1. Curated lists of differentially expressed genes in each stage during human spermatogenesis compared with the curated list of stage-defining genes from Human Atlas.**

| Cell stage | Gene name | Gename from Human Atlas |
|---|---|---|
| Spermatogonia (Sg-1) | "UTF1","C19orf84","BEND4", "FGFR3","LIN7B","MAGEB2", "THRA","MAGEA4","NEIL2","MFHAS1","PAFAH1B3","RGMA", "CDK17","SOX4","HMGA1", "LMO4","TCF3","SNAPC2", "COTL1","SMS","KCNQ2", "TLE1","STK24","DUSP5", "GRN","RFWD3","PIWIL2", "PPP2R1A","LCOR","IRF2BPL" "DAB2IP","CPEB1","USP31" , "ELAVL2","PAFAH1B2","USP11", "DCAF4L1","PHF13", "KMT2B", "TMEM123" , "PABPC4", "PARP1", "CCNI", "POLR2A", "RPSA" , "YWHAB" , "TOMM34", "GATAD2A", "RNPS1", "CELF1", "RPS12" , "RYK" , "RPS19" , "MTPN","MAP4K4" , "TUBA1B", "BANF1", "CBL", "HNRNPDL", "HSP90AB1", "STRN4" , "BCCIP" , "ZNF428" , "ALKBH5", "DYRK1A" , "TKTL1", "RPS5", "CDK2AP1" , "COX7A2L", "CCDC117", "JARID2", "TDRD1","RPLP0", "TRMT112", "CD164", "PHF8", "PDIA4", "NR6A1" , "RPL18A", "RPS21", "PTPA" , "SUMO3", "RAC1" , "RPL38" , | "MAGEA4", "DAZL", "SYCP3", "DMRT1", "DMRTB1", "SOHLH1", "CHEK1" |

| | |
|---|---|
| | "TUBB" , "EEF1B2" , "CBX3", "SPTAN1", "DNAJB6", "BSG", "RPS16", "RPS2" , "RPL18", "RACK1" , "RPS28" , "PFN1" |
| Spermatogonia (Sg-2) | "CT45A6" ,"PNMA5", "SSX3","CT45A10","SSX2","SSX2B", "CT45A9","RHOXF2" , "ZNF280C" , "MCM2","E2F1","MCM6","RHOXF2B","MAGEA4", "CTCFL", "PAFAH1B3","UNG", "HMGA1", "MYBL2", "MCM5","TRMT6", "TLE1","GINS2", "NAE1", "BTG3","TKTL1", "DAZ2","SRM", "DAZ3","DAZ1","PDIA6", "APLP2", "PDIA4", "SLBP","HERC5", "VCX", "CCT6A", "DPEP3", "DMRT1", "NUP93", "XPO1","VCY", "NUDT3", "DAZ4","CENPH", "PRAME", "RNPS1", "ARCN1", "POLR2A", "TSPYL2", "HIST1H4C" "CALR","NOMO2", "DDB1","NFATC2IP" "USP31", "YWHAE", "NASP","ANP32B", "SNRPB","SMC3","RIF1","NOMO3", "NOMO1", "MAP4K4", "PRKDC", "BSG", "PARP1", "RBPJ","CIRBP", "SUZ12", "NCL", "TRMT112","HMGB1", "UBA2","DMRTB1", "SMC1B", "PFN1","HNRNPD","HNRNPDL","MTRNR2L1" "BUD23", "CNOT7", "HSPA5", "PTMA", "HSP90AB1","PABPC4", "MTRNR2L12","MTRNR2L8" |

| | | |
|---|---|---|
| Spermatocyte (Sc-1) | "MAGEA9", "TEX19", "PNMA6E", "JADE3", "MAGEA9B","BEND2", "KIF1A", "MAGEC2", "CLSPN", "CDC6","CTCFL", "ZNF280C","HELLS", "PRIM1", "DPH7","C18orf63" "CCDC73", "CHAF1A", "RAD51AP2" "PRSS50", "VCX3B","DAZ2","DAZ1","GINS2", "TRAFD1", "VCY", "MEIOB", "DAZ3","BTG3","VCY1B", "IQCB1", "VCX", "HIF1A", "ZC3H13", "DAZ4","YTHDF1", "BCAP31", "NPTX2", "VCX3A", "PSMD1", "DPEP3", "SCML1", "TAOK3", "SYCP3", "VCX2","TOP2B", "SAE1","ZCWPW1", "ANKRD31","HIST1H4C" "TEX101", "POLE","TOP2A", "CENPH", "SMC1B", "SFR1", "SMC3","NASP","BUD23", "ABCA5", "PPIG","ACBD3", "HERC5", "CBX1", "SDF2L1","KNL1","HIST1H2BA" "DUT", "SMCHD1", "TSPYL2", "HDAC6", "SUZ12", "BOD1L1", "TKTL1", "ARIH2", "GOLGA2","HORMAD1","NCL", "APLP2", "DNAJC1", "USP9X", "HIST1H2AA" "WEE1","RIF1","PRPF8", "RRM1","U2SURP", "DDX24", "NIPBL", "HMGB1", "DAZL","EIF5B", "SYCP2", "STAG3","SYCP1", "MTRNR2L1", "MTRNR2L12","MTRNR2L8" | "DAZL", "DMRTB1", "SYCP3", "SPO11", "MLH3", "DDX4", "BRCA1", "DMC1", "SYCP1", "SPAG6", "ZPBP2" |
| Spermatocyte (Sc-2) | "LY6K", "C18orf63", | |

| | |
|---|---|
| | "PLEKHG4","DMRTC2", "CCDC172","TRAFD1", "CDCA8", "MEIOB","MDC1", "ANKRD31","ART3", "PIP5K1A","IQCB1","PAN2", "CNTROB", "TEX101", "TEX14","TPTE2","MTF2", "CTSL", "HIST1H2BA" "TPTE", "CCP110", "BUB1", "TOP2B","HORMAD1","NUF2", "SLC25A31","C5orf47","SYCP3" ,"S100PBP","RCN2","PIWIL2","TOP2A","ARIH2","C5orf58","PRC1", "DAZL", "TERF2IP","HMGB2","ARL6IP1", "SYCP1","HNRNPA3","PNISR","TDRD9","SUGP2","TDRD1","NBPF1","DZIP3", "SMC4", "HENMT1", "HSP90B1","SPINT2", "SETX", "KHDRBS1","PRSS21", "TMEM147","SNRPB","HIST1H4C","HSPA2","SENP1","SMC1B","ILF3","BUD23","HNRNPA2B1","SFPQ","LARP1","KDM5B","NBPF3","AC240274.1","RBBP4","PTPA","HNRNPM","HNRNPH3","CCT3", "ILF2", "SRSF11", "VAMP2","MLLT10", "SGO2", "CALM2","SMC3", "CBX3", "KNL1", "KIF5B","CIRBP","NIPBL","HSP90AA1","HIST1H2AA", "HSPA5", "SPATA8" |
| Spermatocyte (Sc-3) | "C9orf57","C4orf46","SLC25A31","SPRTN","TM2D1","CNTROB", "FKBP6", "HIST1H2AA","HIST1H2BA","CKS2", "TERB2","PLK2", "TEX14","CCDC112", "CCP110", |

| | |
|---|---|
| | "CETN3","SLC2A14","FAM174A","PDRG1","TFDP1","COMMD4","PIWIL1","HSPA2","TDRD1"," CTSL", "SLC2A3", "NUF2", "LRRC23","RCN2", "MTX2", "MGAT4D", "SYNGR4", "AC024940.1" "TMEM99", "TYMS", "C5orf58","SELENOS","H2AFZ", "PSENEN", "SPATA8", "SAC3D1", "RFX4", "NT5C3B", "DPY19L2","SPDYA","ZNRF2","SPINT2", "TPTE", "BCAP29", "DYNC2H1","SSNA1","MLLT10", "TERF2IP","TESMIN", "ADAM2","RALGPS2", "ZWILCH", "SENP1","STK31","HMGB2","CALM2","ZMYND10","SETX", "UBN1", "PNISR","ANKRD36C", "EIF5AL1","SNX14","FAM216A", "ANKRD36B", "C15orf48", "KPNB1","ANKRD36","ARL6IP1","KDM5B","HENMT1", "MORF4L1", "SAYSD1", "RBBP4","SUGP2","HORMAD1","KRBOX1", "TUBA3D", "PRSS21", "SBNO1","SPAG6","TPR","NBPF3","AC240274.1" "LYAR", "SPATA22", "NBPF1","TMEM225B", "TUBA3C", "DAZL", "TUBA3E", "SPINK2", "REXO5" |
| Spermatocyte (Sc-4) | "C1orf94","TMPRSS12" "ISOC2","RGCC", "KRT72","GULP1","ADAM2","C |

| | | |
|---|---|---|
| | HST13", "ZBTB32", "AURKA","CCDC86", "CEP55","WDR63","ASH2L","PL A2G6", "ALKAL2", "ZC2HC1C","TBP", "TKTL2","CLGN", "SLC2A5", "CIAPIN1","GLB1L","CCNA1"," TUSC1","C21orf59" "NEK2", "CAVIN3", "C16orf71" "ZPBP2","DAW1", "TYMS", "CCDC42", "LZTFL1", "PCID2","TBPL1", "PPP3R2", "RAE1", "MRPL43", "KPNA5","INO80C", "NUP188", "TCFL5","SNW1", "HSF5","MRPL34", "PBK","REXO5","SPINK2", "MKRN1","ANKLE2", "LYAR", "TDRD7","SNRPC", "PCDH17", "RAN","AHCTF1", "SFMBT1", "COPRS","ASRGL1", "NDUFAF3","ZNF689", "IFT74", "C1orf56","AKAP12", "PTTG1","PPP1R2", "CCNB2","LDHC", "APH1B","TTK","TMEM225B" "WDR87","PLK1", "KDM4D","FBXO25", "SPATA22","FAM216A","KRBOX 1", "ELP5", "CETN1", "ETFRF1", "SPAG6","TYSND1", "ANKRD7", "SMARCA2","C15orf48" "DBF4", "GTF2A2", "CCT6B", "ATP6V1B2" "SPATA16","TUBA3C", "GYG1", "SLC26A8","ZMYND10","SPATA 8", "RFX4", "CENPF", "CCDC110" | |
| Round spermatid (RS-1) | "CTAGE8", "CTAGE4", "CTAGE1", "RGCC", "CDK7", | "ZPBP","DDX4", "SPAG6","ZPBP2", "DNAH6", |

| | | |
|---|---|---|
| | "CDKL2","ISOC2", "WDR63","CEP55","CHST13", "EFHB", "AURKA","NAA20","NBPF11", "CCNB2","NBPF10", "GALNT3", "DNAL4","NBPF19", "C15orf61", "LRRC34", "NBPF12", "NBPF15", "KPNA5","PLK1", "NBPF14", "TTK","CENPF", "CCDC146","PABPC1", "CCDC42","NBPF9","CCDC110","CCDC83", "FBXO25", "MNS1", "HSF5", "NBPF20", "ETFRF1", "C11orf63", "SLC26A8","TBPL1", "PBK","PSMG1","SPATA16","DBF4", "PABPC3", "SNRPC","KRT10", "SMARCA2","WDR87","PTTG1","CAVIN3", "KDM4D","PPP3CC", "TKTL2", "GTF2A2", "HDGFL1", "CCNA1","AC240274.1" "BRD8", "SHCBP1L","ANKRD7", "NBPF3","ZPBP2","CCDC34", "SPAG6","CCDC62", "CLGN", "DYNLT1", "NBPF1","PRRC2C", "DNAJC21","CETN1","IFT74","LDHC", "MRPL34", "ANKLE2", "MRPL43", "LCA5L","SCCPDH", "NEK2", "LYAR", "C16orf71", "CCT6B","TMEM225B", "FMC1", "AKAP12", "REXO5","SPINK2", "CCDC173", "ELP5", "PPP3R2", | "CAMK4", "CREM" |

| | |
|---|---|
| | "COPRS","CNTRL","SPATA22","APH1B","FSIP2", "TUBA3C", "ZPBP" |
| Round spermatid (RS-2) | "C11orf53", "LRRC39", "FAM24A", "LRRC3B", "ENPP2","TMEM243","CCDC81","ODF3L1","OLFML2B","CFAP126","NUTM1","C4orf17","HIPK4","C1orf87", "SOS1", "C11orf97","DEUP1","C4orf47", "CCER1","GOLGA6L2", "CCDC146", "DYNC2LI1", "C1orf105", "LRRIQ3", "EFHB", "ZNF85","USP44","MNS1", "LINC00694","CFAP77", "SSMEM1", "GALNT3", "SPEF2","ADGB", "SPART", "C9orf116", "PPIL6","WDR78","PABPC1", "NBPF15", "NBPF12", "C11orf63", "NBPF10", "ALS2CR12", "NBPF14", "CCDC34", "NBPF20", "C1orf158", "SWT1", "CFAP206","TSSK4","NBPF11", "ZCRB1","DNAL4","CCDC89", "CCDC62", "REEP5","CCNB2","MAATS1", "HIST2H2AA4" "SH3GLB1","CTNNA2", "TBPL1", "ETFRF1","LCA5L","HMGN1","CCDC173","CCDC110","PPP3CC","HIST2H2AA3""DNAJC21","OSCP1","SCCPDH", "PLK1", "PABPC3", "KDELR2", "NAA20","SLC26A8","PBK","GKAP1","C6orf10","ZPBP", "TRIM13","GTF2A2","PPP1R2P3","PSMG1","CCDC42", "FMC1", |

| | |
|---|---|
| | "C7orf61","HSF5", "DBF4","SHCBP1L","FSIP2","LDHC","CCT6B","CETN1","REXO5","ANKRD7","APH1B","LYZL2" |
| Round spermatid (RS-3) | "LRRC3B", "FAM24A", "TPRG1","C4orf17","TJP3","TMEM262","C17orf98", "ODF3L1", "CATSPER1", "C4orf47","EQTN","IGSF10", "C6orf10","PLCH1", "PRSS55","CENPW","CLDN12", "GOLGA6L2","TSSK4","FAIM2", "C1orf105","LYZL4","FAM186A","FAM205A","HIST2H2AA4" "NLRP1","LYZL1","LYZL6", "LYZL2","HIST2H2AA3" "GOLGA6D","SPACA1", "TMEM108","MAP7","SSMEM1", "ATP6V1E2", "MRPL39", "STAT4","LINC00694","ZMYND15","FAM153A","FTMT", "CTNNA2", "GOLGA6A","CCDC82", "POLR1B","SPART","CNTN4","C7orf61","RTKN2","USP44","AFG1L","HMGN1","CCER1","SUN3","MAATS1","DUSP13", "C17orf50", "SNAP29", "RNF148","CCDC89","TFDP2","ACTRT3","FAM153B","DIAPH3","TTLL7","SETD9","TPP2","TEX51","OLAH","ACRV1","SVIP","LRCH4","OSCP1","CFAP206","SPATA46","C2orf16", "FNDC11", "FSIP2","CDKN3","WDR78","CCDC173","CCDC146","KLF5", "ZBTB20", "ZC3H14", "C1orf158", "SPACA3", "CXXC5","BRDT","SPACA4", "MNS1","EIF5B","SPACA7", "CCDC110","PSMG1","FAM209 |

| | | |
|---|---|---|
| | B","FAM209A", "CCDC62", "ACTL7B" | |
| Round spermatid (RS-4) | "CA9","SPACA1", "OLAH", "LRRC52", "SUN5", "EQTN", "CCDC27","TJP3", "FAM205A","RNF148", "CD46", "MAN1A1", "PLCH1","TSSK4", "CD55", "TMC7", "SPACA4", "CRLS1","ACTRT3", "SPAM1","TEX29", "TRIM17","C1orf185","MS4A6E","KLF5","SLC35A5","LATS2","FTMT","SCOC", "FAM153B","ERICH2", "ETNK1","FNDC11", "NDUFB6","LYZL6","C6orf10","SPACA3","LPGAT1","TMEM270","FAM153A","SETD9","BBX","C11orf65","TFDP2","ACRV1","PRR30","ZBTB38", "DGAT2","POLB", "TXNDC2","SAXO1","LYZL2","MEX3C","HINT3","RETREG1","ZC3H14", "FAM209B","SLC38A9","FAM209A","SUN3","C2orf40","TES","DYRK4","LYZL1","SESN3","CCDC168","CALCOCO2","ASB17","ADAM29","LRRC37A2","TEX33","CCDC89","SSMEM1","NDRG3","C2orf42","CPEB2","C20orf173","LYZL4","LRRC37A","HIST2H2AA3","SPATA46","SOX30","CCIN","ACTL7B","HIST2H2AA4" "PRSS55","SERP2","ERGIC2","TEKT5","C7orf61","G2E3","ZBTB20","CXXC5","C8orf88","IZUMO2","SCP2D1","TP53TG5","ZNRF4","CDKN3","AKAP3" | |
| Elongating spermatid (ES-1) | "SLPI","CCDC185", "IQCF6", | "TNP1", "PRM2", "MYO1D", |

| | | |
|---|---|---|
| | "PRSS58","GATA6", "DCDC2C","PRSS37","DNAJB7" , "C10orf120" "TP53TG5", "FAM71A","CCIN","KCNV2", "SRRM5", "CCDC168", "TXNDC2","TEKT5","FAM71B", "FAM57A","CD55","ACTL7B","PCCB","TTLL2", "LTN1", "PCMT1", "FAM209A", "SCP2D1","CA9","CCDC80","ANKRD9","FAM209B", "MS4A6E", "C20orf173" "PDCL2", "GPR18", "ASB17", "G2E3","C2orf40", "FAM8A1","CAST", "TEX29", "RNF151","IQGAP2","CLIP4", "CCDC126","RNF141","C1orf100","SAXO1", "SUN5","GTSF1L","AKAP3", "OTUB2", "ZNF683","MEX3B", "PPP4R1","TRIM42", "SERP2", "ERICH2","MEX3C", "MAPKAPK2","TEX26", "HEMGN", "TSSK2", "C2orf73", "XYLT2", "AKIRIN1", "DXO","LRRC37A2","MARCH8","MKRN2","FBXO39","SPACA4","SUN3","NFKBIB","LRRC37A", "TMEM270", "HMGB4", "TMEM191C","TMCO2", "CHD5","DUSP13","CALCOCO2","RFPL3S","C10orf82","SPPL2C","INPP1", "POLB","CXXC5", "IRGC","SPACA9","C17orf105" "CCDC179", "ZNRF4", "SPACA3","TFAM","TEX37", "SPACA7","ACRV1", "FNDC11","C20orf144" | "TNP1", "HOOK1", "SPATA12", "SPATA18" |
| Elongating spermatid (ES-2) | "TRIM42","CCDC179", "HEMGN", "TTLL2", "BAG5","FAM71B","TFAM", | |

| | |
|---|---|
| | "C1orf100","FAM71F1", "TRIM36","EFCAB1","INPP1", "TUBG1", "KIF2B", "C17orf105","SPACA9","HMGB4","NFKBIB","ACP1","FAM71C", "FAM57A","FBXO39","OTUB2", "COX8C","MARCH8","GOLM1", "TNP2","TEX35", "C3orf30", "PRSS58","TEX37", "IQCF2","TSSK1B","UBE2J1","CCDC196","C10orf82","CCSER2","TLE4","FAM120B","GLRX2","BAG1","IQCF1","MAPKAPK2","SPTY2D1-AS1","LRRD1","TUBA4A","SPATA24", "CA2","HSPA1L", "CDC14A","SPERT", "KNSTRN","DNAJA4","FAM71E1", "AKAP4", "AKIRIN1", "GSTO2", "TNP1","C2orf88", "FAM81B","RNF151","IRGC","FNDC8","GNG2","CAPZA3","AC010255.3","TEX44","ANKEF1","CEP170","FAM71A","CNN1","AL672043.1","TSSK2","CKB","HOOK1","GAPDHS","OXCT2","GTSF1L","CT83","PRM1","RNF141","SPATA6","RFPL3S","CXCL16", "OAZ3","TSSK6", "TMCO2", "SPATA18", "SH3RF2","CABS1", "AKAP1","LELP1", "TPPP2", "PRM2","GLUL","CRISP2","C20orf141","ODF1","TMEM31","SAMD4A" |
| Elongating spermatid (ES-3) | "PRM1","LEMD1","TNP1","AC010255.3","SPATA19","C9orf24","IQCF3","TNP2","AL672043.1","SPEM1","SPATA6","MORN3","TAF10","PRM2","CRISP2","SMCP","TSSK6","SPATA20","ODF1","PCP2","SH3RF2","H1FNT","IQCF |

| | |
|---|---|
| | 1","LELP1","ODF2","PSMF1","FNDC8","SPATA32","PHF7","DYNLL2","ESS2","AKAP1","ARPP19","TSPAN16","CYLC1","C17orf74","ACSBG2","RNF138","CLMN","WDR1","FUNDC2","C11orf71","PAQR7","DCUN1D1","CXCL16","TUBA4A","KIF2C","CAPZA3","TSPAN1", "CA2","C19orf70","SOCS7","NUPR2","IQCF2","TMEM31","OAZ3","TSPAN6","NRDC","GAPDHS","CLPB","TEX44","FSCN3","SPATA3","DCAF1","CCDC91","C10orf62","LPIN1","TPPP2","ADIG","GLUL","LINC00854","ACTL7A","C16orf82","AKAP4","UBE2J1","ADRM1","NSUN4","ACAP1","ETNK2","MFAP3L","BAG1","HOOK1","FAM81B","CABS1","CEP170","HSPA1L","ACTRT2","CCSER2","ISG20L2","C12orf54","CCDC54","SPERT","ACE","GPX4","SAMD4A","SPATA18","TEX37","C20orf141","SPTY2D1-AS1","PRH1" |
| Elongating spermatid (ES-4) | "C16orf78", "SPATA3", "GSG1", "PHOSPHO1", "C10orf62", "FSCN3","PRM2", "PROCA1", "CRISP2", "LELP1","MOSPD3", "AC133555.3" "GLUL", "RANGAP1", "DCUN1D1","C16orf82", "MORN3","LPIN1","ACAP1","PAQR7","TCP11", "PCYT2","CLPB", "RNF138", "ACSBG2", "SPATA18","BPIFA3", "FAM46C","GAPDHS","DDX3X","ZFAND3","SMKR1","TSSK3","ODF1", "NSUN4", "BOD1L2", |

| | |
|---|---|
| | "TSSK6","SMCP","MROH7","CABS1","CXCL16","RND2","DNAJC4", "AC106782.1" "OAZ3", "C3orf22","MS4A14", "C17orf74", "ODF3L2", "REEP6","NUPR2","ETNK2","CCNY", "TSPAN6", "HIP1", "ABHD1","NDUFA13","CCDC91","SPATC1","TPPP2","STPG3","TSPAN16","FAM71F2","FUNDC2","ZDHHC19","UBE2N","SPEM1","RCC1", "PCP2", "RNF44", "MFAP3L","TTC7A","TEX46","PKM","C19orf70", "AZIN2","METAP1", "AKAP1","DNAJB8", "LINC00854","LRRD1","DCAF1","AKAP4","CA2", "DGCR6L", "WBP2NL", "ODF2", "SH3RF2", "RAB11FIP4","ACTL7A", "PARD6A","ISG20L2","GPX4", "TPI1", "AL672043.1" "C12orf54", "UBQLNL", "STARD10", "MEX3D","PRM1" |

### Supplementary Table 2

#### Table 2. Lists of regulon markers from each individual stage during human spermatogenesis.

| Stages | Marker regulons |
|---|---|
| Sg-1 | "RUNX3","BARHL2","ZNF362","CDX2", "IRX2", "POLR3G","NFATC1","NFIC", "CLOCK","PHF21A","SIX1", "NR2C1","SOX8", "HMGA1","MLXIP", "NR1H2","FOXA1","SMC3", "ESX1", "LHX3", "RFX7", "CTCFL","SOX4", "POU3F1","E2F6", "YY2",,"PAX8", "POLR3A","HMGB1","E2F1", "MAZ",,"IRF3","SMARCC2","ZNF319","FOXK1","JUN",,"TCF12","NR3C1","HSF2 ", "KLF4", "MBD1", "ZNF646","FOXP1","GMEB1","ZNF233" |
| Sg-2 | "GLIS3", "POLR3G","NFIC","PHF21A","SIX1","DDIT3", "SOX8","GMEB1", "NR2C1", "HMGA1", "IRX2","SMC3","FOXD2", "ATF6B", "CLOCK", "CTCFL", "ESX1","PAX8","HMGB1", "TCF12", "E2F1","POLR3A","E2F6","SOX4","JUN", "POU3F1","MAZ", "YY2", "SMARCC2" "IRF3", "ZNF319","HSF2","NR3C1", "ZNF646","MYBL1" |
| Sc-1 | "PITX2", "PAX6","TLX2","ATF6B", "ETV6","ZNF579","FOXF2", "CLOCK", "GMEB1", "FOXK1", "POLR3G","RXRB","FOXD2", "HMGA1", "SMC3", "PHF21A","CTCFL","MECOM","ESX1","PAX8","NFATC1","HSF2","IRF3","SMAR CC2" "NR2C1", "E2F1","MAZ", "HMGB1", "ZNF319","POLR3A","YY2", "E2F6","TCF12", "MYBL1", "NR1H2", "JUN", "NR3C1", "ZNF646" |
| Sc-2 | "RXRB","DDIT3", "GMEB1", "POLR3G","FOXD2", "PHF21A","ATF6B", "CLOCK", "SMC3","CTCFL", "PAX8","IRF3","MYBL1", "ZNF319","MAZ", "SMARCC2" "TCF12", "HMGB1", "POLR3A","E2F1","YY2","HSF2" |
| Sc-3 | "LEF1","FOXD2","ZNF546","CLOCK","TCF12","MYBL1","IRF3","MBD1","SMC3", "NR1H2","OVOL1" |
| Sc-4 | "DTL","ZFP64","OVOL1","MBD1","SOX18","MYBL1","ZNF233","ZNF646","KDM4 D","TCF12" |
| RS-1 | "DTL","ZFP64","OVOL1","MBD1","SOX18","MYBL1","ZNF233","ZNF646","KDM4 D","TCF12" |
| RS-2 | "KDM4D","SOX18", "ZNF646", "HSF2", "ZNF233" |
| RS-3 | "HSF2", "KDM4D","BBX","NR3C1","ZNF646" |
| RS-4 | "BBX","GFI1B","NR3C1" |
| ES-1 | "SHOX2","GATA6","BBX","GFI1B","KLF4" |

| | |
|---|---|
| ES-2 | "GATA6","DUX4","PRKAA1","GFI1B","KLF4" |
| ES-3 | "DUX4","PRKAA1","BCL6","FOXP1" |
| ES-4 | "TAL2","NFE4","FOXB1","PRKAA1","DUX4","ZNF85" |

### Supplementary Table 3

**Table 3. Lists of genes involved in DNA repair pathways consisted of Nuclear excision repair (NER), Base excision repair (BER), Mismatch repair (MMR), Homologous recombination (HR), Non-homologous end joining (NHEJ), and Inter-crosslinking repair (ICL).**

| Pathway | Gene name | Reference |
|---|---|---|
| Nuclear excision repair (NER) | "ERCC4","XRCC4","ERCC6", "TCEA1", "ERCC8", "XPC", "ERCC1", "POLR2A", "STK19", "RFWD3", "ELOF1", "ERCC2", "UVSSA", "XPA", "GTF2H4", "ERCC5", "ERCC3", "POLK", "GTF2H1", "GTF2H5", "RFWD2", "REV3L","MMS19" | (Boeing et al., 2016; Calmels et al., 2016; Cheng et al., 2000; Friboulet et al., 2013; Gayarre et al., 2016; Joo et al., 2016; Kaina, 2020; Kou et al., 2008; Lin et al., 2018; Lipkowitz and Weissman, 2011; Lisica et al., 2016; Ma et al., 1994; Manandhar et al., 2015; Nakazawa et al., 2012; Ogi and Lehmann, 2006; Petruseva et al., 2014; Ræder et al., 2018; Rosin et al., 2015; Soltys et al., 2013; Song et al., 2017; Sugitani et al., 2016; Yang et al., 2018) |
| Base excision repair (BER) | "NTHL1", "LIG1", "TDP1", "PARP1", "XRCC1","REV1" | (Limpose et al., 2018; Nazarkina et al., 2007; Prasad et al., 2016; Rechkunova et al., 2015; Sukhanova et al., 2010; Sykora et al., 2013) |
| Mismatch repair (MMR) | "MSH2", "MLH1", "MSH6", "PMS2" | (Ellison et al., 2004; Kantelinen et al., 2012; Nakagawa et al., 2004) |
| Homologous recombination | "FIGNL1", "RMI1", "BLM", | (Amendola et al., 2017; Boudrez |

| | | |
|---|---|---|
| (HR) | "RMI2", "MCM8", "HROB", "MCM9", "RAD51", "RAD51C", "XRCC3", "BRCA2", "PALB2", "BRCA1","RAD51B", "NBN", "BARD1", "RAD51D", "MRE11A", "XRCC2", "SFR1", "EME1", "RAD54L", "AUNIP", "MUS81", "TRAIP", "FEN1", "SCC1", "DDX11", "RNF8", "XRCC1", "RNASEH2C", "RNASEH2A", "LMNA", "SCAP", "MMGT1", "UFSP2", "WDR83", "WDHD1", "ECHS1", "C7orf26","TCEB2", "PPP1R8", "DHX9","SLC25A28", "PDS5B", "HUWE1", "MAU2", "CDK2", "PPP4C", "FOXM1", "CDC25", "ESCO2", "KDM8", "STAG1", "JMJD6", "MED12", "DIS3", "DKC1", "TIPRL", "DBF4" | et al., 2000; Brenneman et al., 2002; Brommage et al., 2014; Buisson and Masson, 2012; Chon et al., 2009; Couturier et al., 2016; Domingo-Prim et al., 2019; Ertl et al., 2017; Froyen et al., 2012; Gannavaram et al., 2014; Gaponova et al., 2017; Gu et al., 2008; Hanada et al., 2006; Heeke et al., 2018; Hickson and Mankouri, 2011; Hiller et al., 2012; Holloman, 2011; Huo et al., 2020; Hustedt et al., 2019; Johnson et al., 1999; Junes-Gill et al., 2014; Kikuchi et al., 2005; Konstantinopoulos et al., 2015; Lee and Pelletier, 2016; Lou et al., 2017; Lu et al., 2012; Ma et al., 2017; Matos et al., 2008; Mondesert et al., 2002; Moon et al., 2012; Nishimura et al., 2012; Pal et al., 2017; Park and Lee, 2020; Redwood et al., 2011; Reh et al., 2017; Singh et al., 2008; Soo Lee et al., 2016; Su et al., 2012; Su et al., 2016; Takata et al., 2000; Tan et al., 2007; van der Lelij et al., 2009; van Schie et al., 2020; Viera et al., 2009; Villoria et al., 2019; Vispé et al., 1998; Walter et al., 1996; Wang and Wang, 2014; Wilson et al., 2012; Wu and Yu, 2012; Wu et al., 2012; Yang et al., 2016; Zha et al., 2009; Zhang, 2013; Zhang et al., 2020; Zhao et al., 2017) |
| Non-homologous end joining (NHEJ) | "C7orf49", "ERCC6L2", "POLL", "UIMC1", "IREB2", "SHLD2", "POLQ","SHLD1", "DCLRE1C", "RIF1" | (Chapman et al., 2013; Chifman et al., 2014; Dev et al., 2018; Dianatpour and Ghafouri-Fard, 2017; Felgentreff et al., 2015; Francica et al., 2020; Schimmel |

| | | |
|---|---|---|
| | | et al., 2017; Slavoff et al., 2014; Waters et al., 2014) |
| Inter-crosslinking repair (ICL) | "ESCO1", "FANCD2", "SLX4", "USP1", "DCLRE1A", "FANCG", "C1orf86","C17orf70","FANCM" , "FANCB", "FANCA", "FANCF", "C19orf40", "FANCL", "FANCC", "FANCI", "TRAIP", "UBE2T","BRIP1" | (Andreassen and Ren, 2009; Deans and West, 2011; Gunn et al., 2016; Huang et al., 2019; Jiang et al., 2017; Kim et al., 2012; Leman and Noguchi, 2014; Ling et al., 2007; Liu et al., 2010; Murai et al., 2011; Wu et al., 2019; Yamamoto et al., 2011) |
